# Unravelling traits with complex inheritance mode: a gene panel perspective

**DOI:** 10.1101/2024.12.14.628476

**Authors:** Joanna Szyda, Tomasz Suchocki, Monika Gołąb, Bartłomiej Budny, Magda Mielczarek, Jakub Liu, Krzysztof Kotlarz, Elżbieta Wrotkowska, Marek Ruchała, Katarzyna Ziemnicka

## Abstract

Although gene panels for complex phenotypes target genes contributing significantly to phenotypic variation, identifying causal mutations remains challenging. We aimed to define the process of analysing gene panel data to uncover genomic insights for traits with complex inheritance. The implemented procedure included the exploration of (i) the association between phenotypic variability and the variability of SNPs within genes, (ii) the association attributed to specific variants, (iii) causality potential by analysing the genomic context in the vicinity of the significantly associated SNPs (iii_a) either through clustering – to identify variants with genotypic patterns similar to the significant SNPs, or (iii_b) through smoothing the set of original P values to incorporate the information on the impact of the particular mutations, as expressed by the SNP-specific PhRED-CADD scores. This procedure was applied to the data from the obesity gene panel. No significant gene-level associations were found. However, the SNP-level results were promising in narrowing down the initially available set of variants to determine the true causal mutations. We introduced a procedure for analysing gene panel data to identify potentially causal mutations for phenotypes with complex inheritance patterns. The procedure allows for narrowing down the originally available set of variants offered by a gene panel, consisting of genes important for the phenotype under study, to SNPs that have a statistically and genomically validated potential to be causal in modifying protein structure.

## 1. Introduction

The importance of next-generation sequencing in medical genetics cannot be overestimated. It allows for the identification of polymorphic variants and testing of their association with clinical phenotypes or, for known associations, for predicting disease risk (Dahui, 2019). Polymorphisms can be identified based on panel, whole-exome (WES), or whole-genome sequencing (WGS) data, listed in the order of increasing scope of polymorphism number. Although WES and WGS provide a comprehensive way to describe genomic variation and thus enable much broader genetic insight into the genetic architecture of phenotypes, especially phenotypes underlying a complex mode of inheritance, gene panels also offer features that make them important for clinical reasons. The major advantage of gene panels is that they are much more cost-effective than WES and WGS, making them accessible to a large number of individuals. Not only because they are a much cheaper diagnostic option than WES and WGS, but also because they provide a faster turnaround time. Furthermore, the reduced volume of data does not require large amounts of resources and simplifies the technical aspects of raw data processing and downstream bioinformatic analysis (Resta et al., 2018). Faster sequencing and analysis compared to WES and WGS is an essential factor in clinical scenarios where faster diagnosis impacts patient management and treatment (Kryukov et al., 2009).

However, although sequencing a predefined list of genes associated with specific diseases can provide a more relevant diagnosis, especially when clinical suspicion is strong for a particular condition (Resta et al., 2018; Quaio et al., 2021; Rehder et al., 2021), it remains less straightforward to analyze disorders with an underlying complex mode of inheritance. Since such phenotypes are typically determined by many genes with various effects magnitudes, the diagnosis and risk assessment based on gene panel data are less accurate than when based on WES and WGS. On the other hand, for such disorders, one wants to profit from the accessibility of gene panels that allow for the ascertainment of large cohorts covering a broader variety of non-genetic effects and thus allowing for a more accurate estimation of genetic effects as well as allowing for achieving satisfactory power of testing, that for a complex trait is often a strongly limiting factor (Kryukov et al., 2009) precluding the accurate assessment of disease risk associated with particular variants.

Panels properly designed for a given complex disorder are composed of genes responsible for a significant proportion of the observed phenotypic variation. However, it often remains difficult to select which available variants represent the true causal mutation. Therefore, the objective of our study was to explore the pathway to extract genomic information from gene panel data downstream of variant identification in the context of traits underlying a complex mode of inheritance. The gene panel data for 190 women with BMI records was used to illustrate the analytical procedure.

## 2. Materials and Methods

### 2.1. The cohort

The analysis pipeline was exemplified using a cohort of 190 adult, unrelated women (age range 23 to 45 years), unrelated women with a Body Mass Index (BMI) ranging from 17.50 to 87.00 with an average BMI of 29.88±10.41. Each individual was genotyped by the custom NGS panel consisting of 22 hormone receptor genes located on fourteen chromosomes (1, 3-8, 11, 12, 14, 15, 17-19) that were identified in the literature as important for regulation of metabolism and obesity: adiponectin receptor 1 (*ADIPOR1*), adiponectin receptor 2 (*ADIPOR2*), adrenoreceptor β 3 (*ADRB3*), apelin receptor (*APLNR*), arginine vasopressin receptor 1B (*AVPR1B*), caveolin 3 (*CAV3*), epiregulin (*EREG*), estrogen receptor 1 (*ESR1*), estrogen receptor 2 (*ESR2)*, fibroblast growth factor receptor (*1FGFR1*), ghrelin receptor (growth hormone secretagogue receptor, *GHSR*), G protein-coupled estrogen receptor 1 (*GPER1*), insulin like growth factor 1 receptor (*IGF1R*), insulin receptor (*INSR*), leptin receptor (*LEPR*), melanocortin 2 receptor (*MC2R*), visfatin receptor (nicotinamide phosphoribosyltransferase, *NAMPT*), glucocorticoid receptor (nuclear receptor subfamily 3 group C member 1, *NR3C1*), aldosterone receptor (nuclear receptor subfamily 3 group C member 2, *NR3C2*), thyroid hormone receptor α (*THRA*), thyroid hormone receptor β (*THRB*), and vitamin D receptor (*VDR*). This panel was designed using an algorithm developed by Thermo Fisher Scientific Ampliseq Designer (http://www.ampliseq.com).

For genotyping, DNA was extracted from the patients’ peripheral blood at the Molecular Endocrinology Laboratory, Department of Endocrinology, Metabolism and Internal Diseases at Poznan University of Medical Sciences. The Ion Torrent Personal Genome Machine (Ion PGM)™ (Thermo Fisher Scientific) was used for the analysis. The construction of the library and subsequent enrichment of the paired DNA samples was conducted using the Ion OneTouch v2 system according to the manufacturer’s protocol (Thermo Fisher Scientific). The Ion PGM Sequencing 400 Kit reagents and Ion 316 v2 sequencing chips were used for sequencing. The average coverage was 199x. The quality filtering of the reads was performed using Ion Reporter software (Thermo Fisher Scientific) and Single Nucleotide Polymorphisms (SNPs) were called compared to the human reference genome GRCh37 / hg19. The panel revealed 262 SNPs, and the number of SNPs varied considerably between genes, ranging from just a single SNP within THRA to 40 SNPs within INSR (Fig. 1). The within-gene pairwise linkage disequilibrium pattern between SNPs, expressed by the R^2^ statistics, was visualised on Fig. 2.

**Fig. 1.**
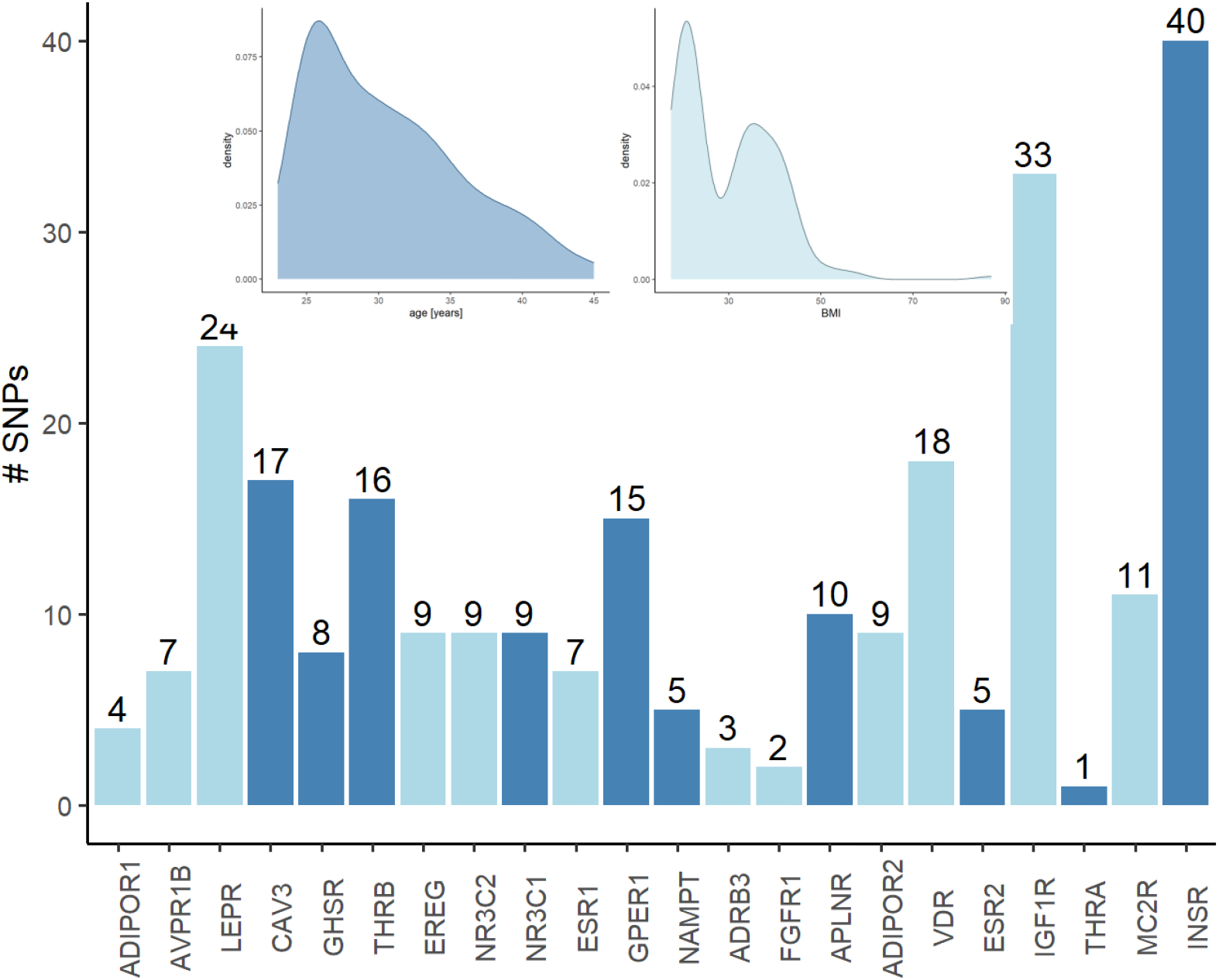
The number of SNPs within each gene covered by the panel and smoothed age and BMI frequencies in the sample of 190 women.

**Fig. 2.**
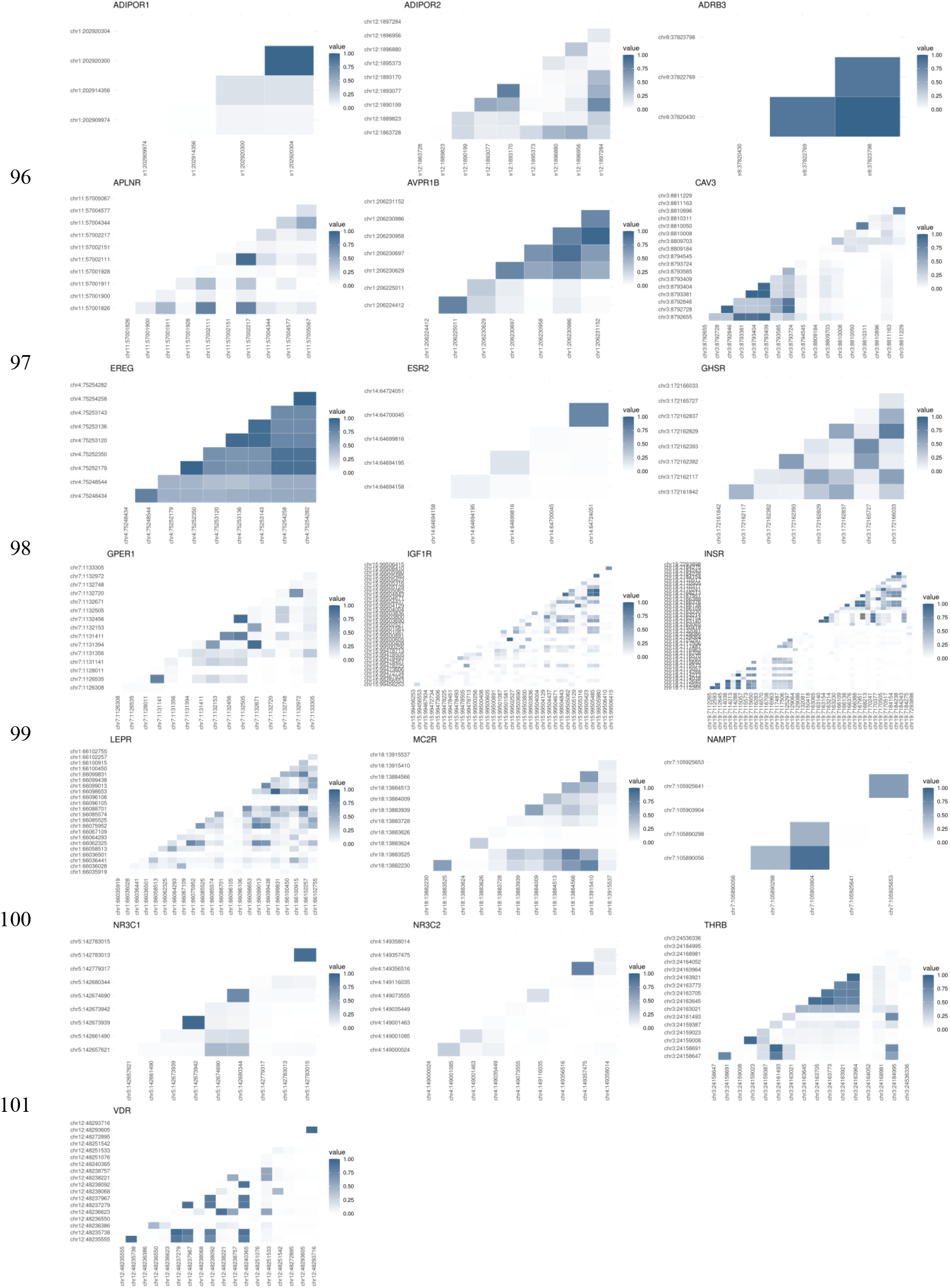
Structure of linkage disequilibrium within each gene expressed by R2 between pairs of SNPs.

### 2.2. Modelling gene effects cumulated over all SNPs

The sequence kernel association weighted variance component test (SKAT) was used to test the general significance of all SNPs representing a given gene (Wu et al., 2011). The test statistic is given by:

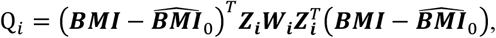

where ***BMI*** is the vector of observed BMI values for the 190 individuals, 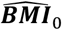 is the vector of BMI predicted by the null model without the genetic component, ***Z***_***i***_ is the SNP genotype matrix for gene *i* which was parameterized as 0, 1, or 2 for a homozygous, heterozygous and an alternative homozygous SNP genotype, respectively, and ***W***_***i***_ is the diagonal matrix of weights for each SNP within gene *i*, that were represented by PhRED-scaled CADD scores for each SNP. Note that to avoid the removal of genotypic variation underlying SNPs with zero CADD scores during matrix multiplication, a small number *ε=0.01* was added to each PhRED-scaled CADD score that was equal to zero.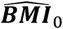 was predicted based on the OLS (Ordinary Least Squares) estimates of 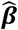 from the following null model ***BMI*** = ***μ*** + ***Xβ*** + ***e***, where ***β*** comprised the effects of: height, weight, waist circumference, hip circumference, age at data ascertainment, blood glucose level, blood insulin level and blood insulin level 120 min after a meal, blood High-Density Lipoprotein level, Homeostasis Model Assessment index, and Quantitative Insulin Sensitivity Check index, ***e*** represents the error term and ***X*** is a design matrix linking individuals to effects from ***β***. The asymptotic distribution of the SKAT test under the null hypothesis follows a mixture of 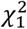 distributions for which the nominal P values can be approximated following (Davies, 1980). The SKAT test was implemented via the R package SKAT 2.2.5 (10.32614/CRAN.package.SKAT).

Genomic covariances between individuals were quantified based on their identity-by-state SNP genotype similarities, calculated separately within each gene and expressed as the genomic ***r***elationship matrices (***G***_***i***_) following (VanRaden, 2008), where 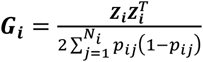 with the subscript *i* corresponding to a gene, subscript *j* to a SNP, ***N***_***i***_ representing the number of SNPs in the *i*-th gene and 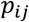 is the SNP minor allele frequency. Genomic relationship matrices were estimated using the R package AGHmatrix 2.1.4 (10.32614/CRAN.package.AGHmatrix).

### 2.3. Modelling effects of individual SNPs

The association study was carried out by fitting the following mixed linear model simultaneously to all SNPs in the panel:

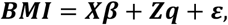

***w***here vectors of ***BMI*** and ***β*** are as defined above for SKAT, ***q*** is a vector of random SNP effects and ***ε*** represents the vector of random residuals, ***Z*** is a SNP genotype matrix for all SNPs as defined above. The covariance structure of the model is given by 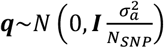 and 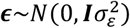, where ***I*** represents an identity matrix, 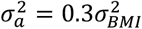 is the additive polygenic ***v***ariance, ***N***_*s****N****P*_ = *262* is the number of SNPs, and 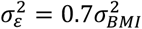 represents the residual ***v***ariance. Consequently, the variance of ***BMI*** is given by: *var****(BMI)*** = ***ZGZ***^***T***^ + ***R***, where 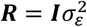 and 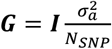. The estimation of model effects was based on solving the mixed ***m***odel equations introduced by (Henderson, 1984) using a custom-written R script.

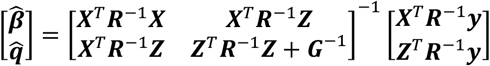

Note that the variance components 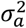 and 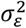 were not estimated in this study but were assumed. For testing the hypotheses: ∀*i* ∈ {*1*, …, *N*_*sNP*_} *H*_*0*_: *q*_*i*_ = *0* vs *H*_*1*_: *q*_*i*_ *≠ 0*, the Wald test 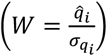 was used, where 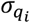 is the standard error of the estimated SNP effect *q*_*i*_. Under H_0,_ ***t***his statistic asymptotically follows the standard normal distribution.

Furthermore, hierarchical clustering of SNP genotypic variation was used to explore genetic (dis)similarity between SNPs to validate whether the significant variants from the above model represent a larger group of SNPs or only express individual associations. Clusters were identified using the iclust function in the R package psych 2.4.6.26 (10.32614/CRAN.package.psych). The last step of the individual SNP analysis consisted of prioritizing functionally important SNPs by smoothing the -log10 transformed P values from the mixed linear model applied by calculating the average P value over each pair of neighbouring SNPs weighted by their PhRED-scaled CADD scores using a custom written R script.

## 3. Results

### 3.1. Gene-based results

No significant association was detected between the genotypic variation expressed by all SNPs within each gene and the variation in BMI. None of the P values of the SKAT test favoured the genomic over the null model, with the lowest nominal type I error being 0.10499 for *EREG* (Table A1). Also, the genomic relationship matrices, expressed as heat maps for visual interpretation, did not reveal any pattern of (dis)similarities between individuals with concordant BMI phenotypes (Fig. A1).

### 3.2. SNP based results

On the other hand, when the association of individual SNPs was estimated with the linear mixed model for ten SNPs located within five genes, a significant additive effect on BMI was estimated (Table 1). The only exonic SNP was a missense variant rs11544331 located in *GPER1* (P=0.01872), representing a C to T substitution with the PhRED-scaled CADD score of 5.85. *IGF1R* harboured the largest number of four significant SNPs spanning 27267 bp, with significance varying between P=0.00499 (rs2293117) and P= 0.04695 (rs2684786) that are located either in introns or in the 3’UTR of the gene. In contrast, the significant (P=0.02571) intronic rs2684785 had the highest PhRED-scaled CADD score of all the significant SNP amounting to 14.37. *INSR* and *VDR* each contain two very closely linked significant SNPs located in introns 276 pb (rs2860178, rs7251963) and 111 bp (rs11168293, rs11168292) away from each other, respectively. With P= 0.00319, rs11168293 from *VDR* was the most significant variant. In *THRB*, a significant (P= 0.01276) variant is located in 3’UTR.

**Table 1.**
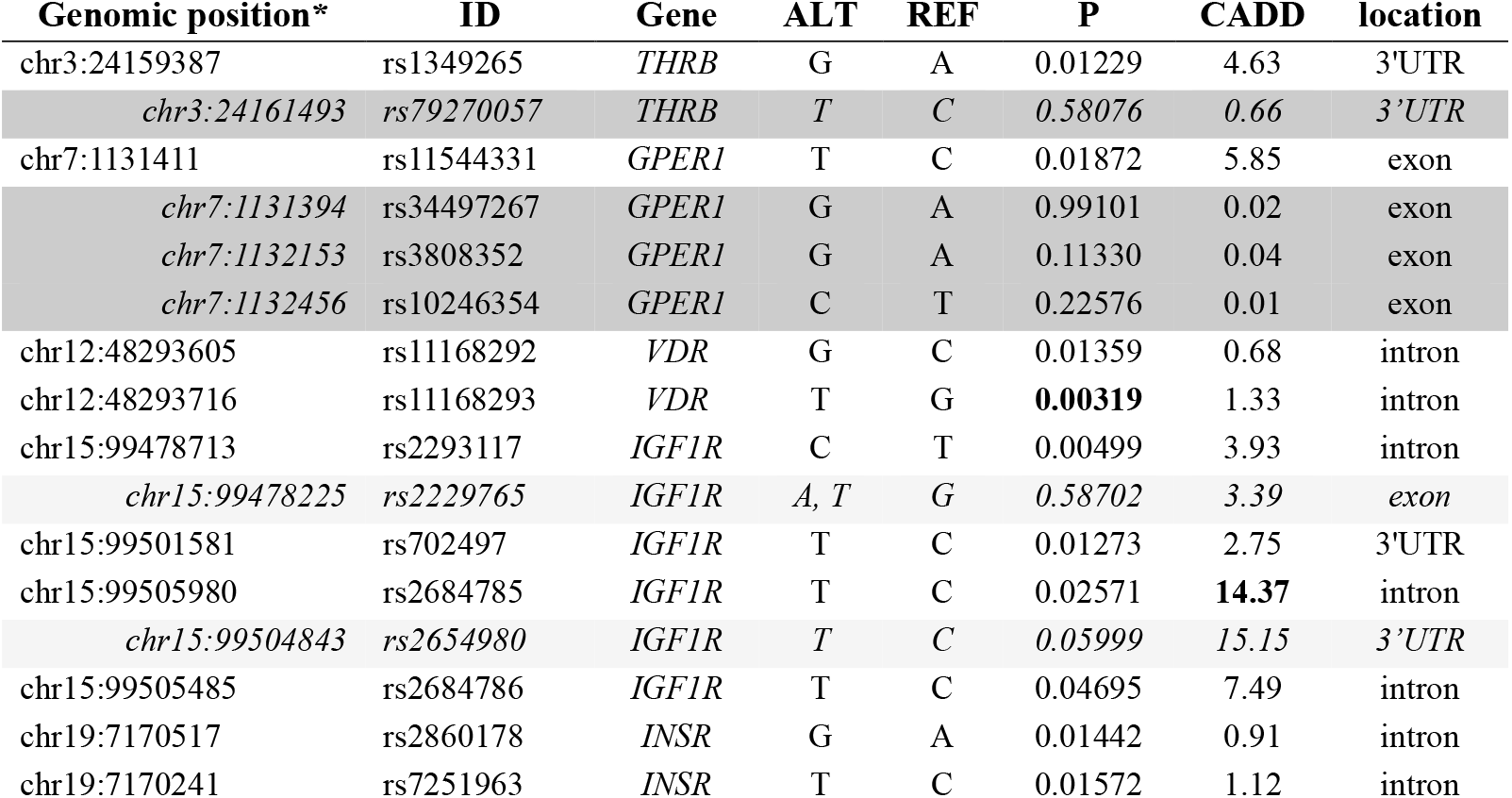
Significant SNPs from the linear mixed model (no shading) complemented by SNPs identified by the hierarchical clustering approach (light shading) and the P value smoothing approach (dark shading).

Exploring SNP genotype variability within genes marked by significant SNPs using hierarchical clustering revealed that for *THRB* and *GPER1*, the genotypic variation of the most significant variant did not directly cluster with the surrounding SNPs. Both significant variants in *VDR* and *INSR* formed a common cluster with each other and did not include other variants. A more complex pattern emerged for the four significant SNPs located within *IGF1R*. The intron variant rs2684786 and the 3’UTR variant rs702497 did not cluster with other SNPs. However, rs2293117, located in an intron, directly clustered with an exon variant rs2229765, while another intron variant rs2684785 directly clustered with rs2654980 located within 3’UTR but characterized by a high PhRED-scaled CADD score of 15.15. Thus, both SNPs can also be considered potential causal candidates for modifying the risk of obesity (Fig. 3, Table 1).

**Fig. 3.**
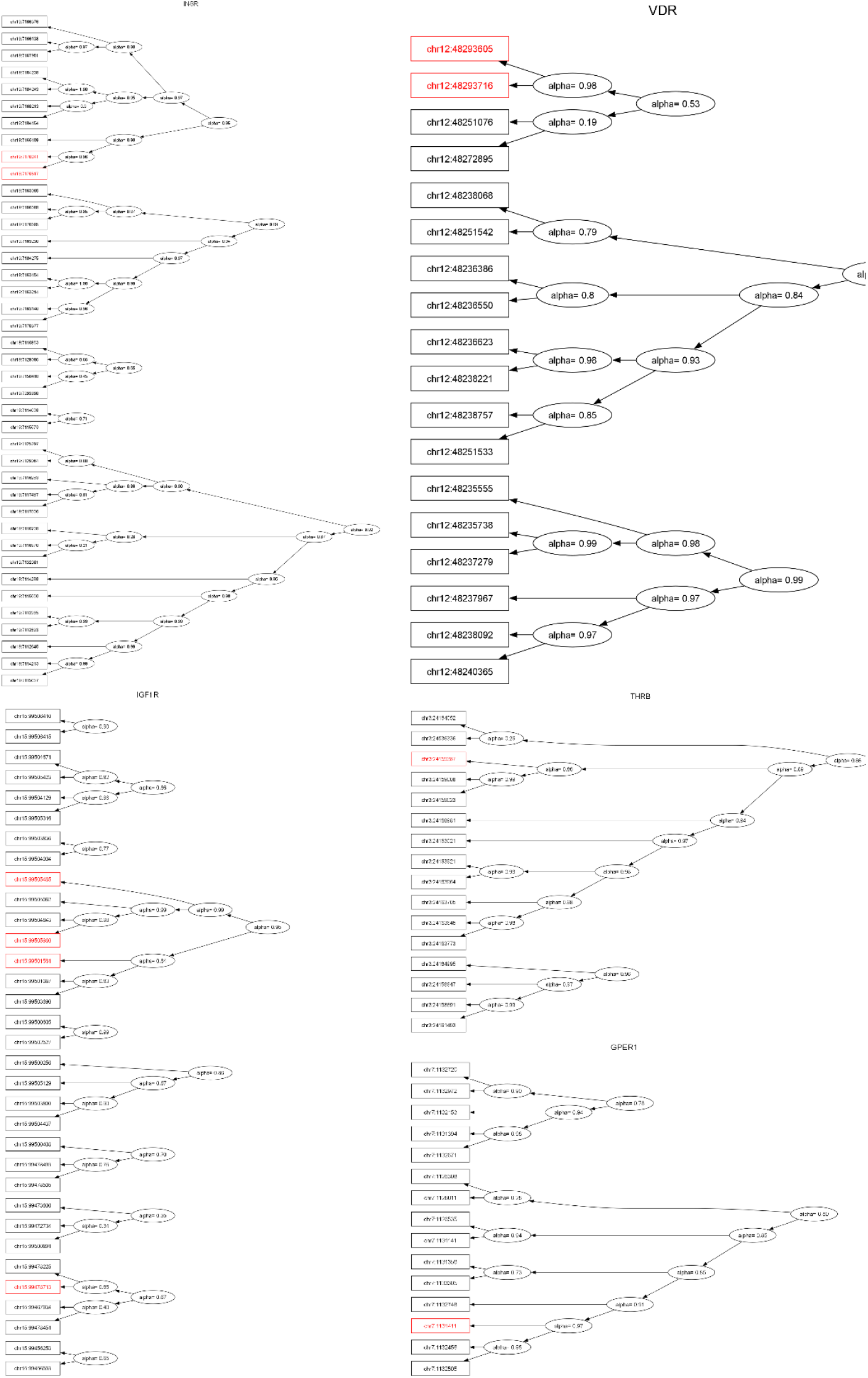
The hierarchical clustering of SNPs within genes significant in the linear mixed models that are marked in red. Alpha represents the ratio of genotype variance expressed by particular SNPs and variance expressed by the cluster.

The final aspect of exploring the interplay between significant SNPs and other SNPs located within the given gene was SNP prioritization based on their nominal P values from the linear mixed model, position within the gene and their functional importance expressed by PhRED-scaled CADD score. The results of P value smoothing, presented in Fig. 4 indicated an additional significant SNP from *THRB* and three additional significant SNPs from *GPER1* (Table 1).

**Fig. 4.**
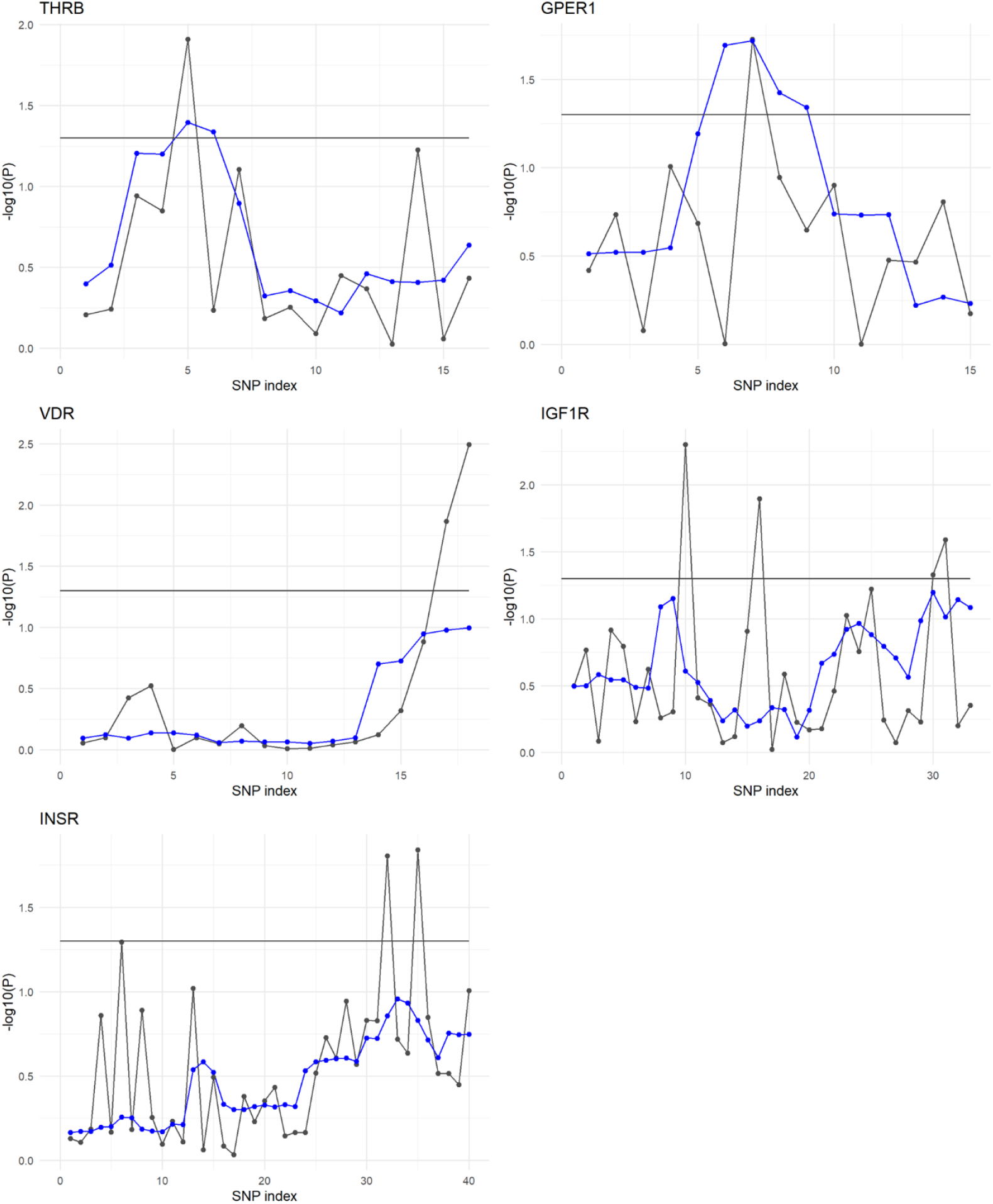
Original -log_10_(P) from the linear mixed model (black dots) and -log_10_(P) smoothed by averaging P values over 3 SNPs weighted by their PhRED-scaled CADD scores (blue dots).

## 4. Discussion

In this study, we implemented a three-step approach that starts with (i) exploration of the association between phenotypic variability and the variability of SNPs within genes. Then, we moved to (ii) exploration of the association attributed to specific variants. Furthermore, anchoring on SNPs significant in the association analysis, (iii) we explored the potential of causality by analysing genomic context in the vicinity of the significantly associated SNPs (iii_a) either through clustering – to identify variants with genotypic patterns similar to the significant SNPs, or (iii_b) through smoothing the set of original P values to incorporate the information on the impact of the particular mutations, as expressed by the SNP-specific PhRED-CADD scores (Fig. 5).

**Fig. 5.**
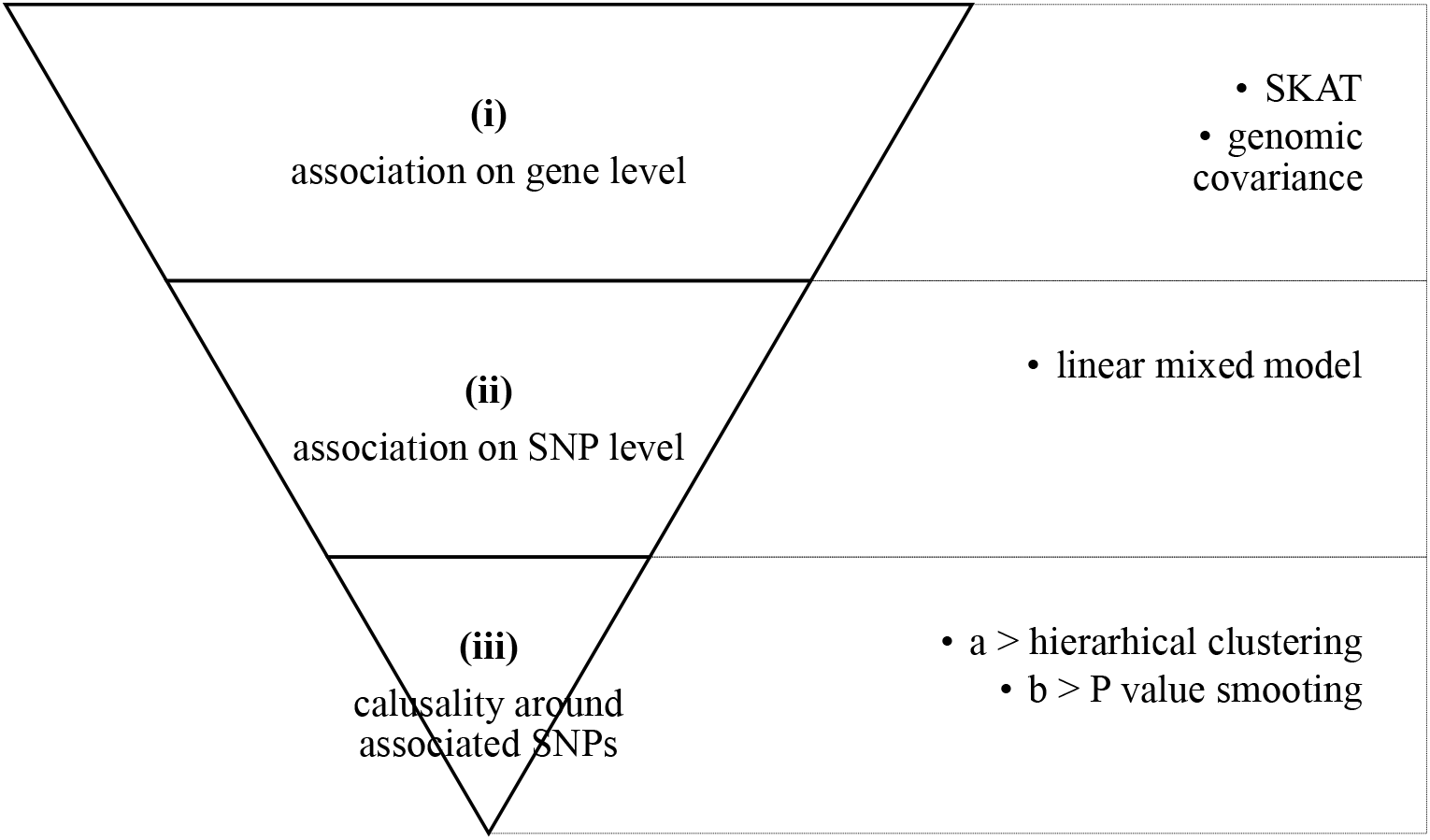
The block diagram of the pipeline for the sequential exploration of gene panel data for a trait with a complex mode of inheritance.

The above procedure was applied to the gene panel data for obesity. No significant association was observed on the gene level. Then, by considering the association of particular SNPs estimated by the linear mixed model, the following significant variants were concordant with the literature. rs2293117, rs2684786, and rs2684785, located in an intron of *IGF1R*, were listed in the ClinVar database as associated with the risk of growth delay due to insulin-like growth factor I resistance. Moreover, rs702497 from the 3’UTR of *IGF1R* was reported as responsible for the variation of waist-to-hip ratio adjusted for BMI (Zhu et al., 2020). It was also listed in the ClinVar database as associated with the risk of growth delay due to insulin-like growth factor I resistance. In *INSR*, the significant intron variant rs7251963 was reported as significant for body height in the large-scale GWAS of 5.4 million individuals (Yengo et al., 2022). Since none of the abovementioned SNPs represents a functional mutation with a possibility to directly alter a protein sequence, we then focused on exploring the variants in proximity of those significant SNPs. Among them, the hierarchical clustering approach indicated rs2229765 that is located in the exon of *IGF1R* and was listed in the ClinVar database as associated with the risk of growth delay due to insulin-like growth factor I resistance as well as described as a modifier of the weight response to a lifestyle change intervention (Martínez and Milagro, 2015). Another, albeit non-exonic variant of *IGF1R*, rs2654980, was reported to be associated with BMI in the EBI GWAS catalogue (study ID GCST90255621). The approach of smoothing P values based on their functional importance expressed by the PhRED-scaled CADD scores highlighted 3 exonic variants of *GPER1* (rs34497267, rs3808352, rs10246354). Still, none of them were reported in the literature as being related to obesity. Providing the very small sample size analysed in our study, further evaluation of those variants in a much larger cohort would resolve their potential causal effect on the risk of obesity.

## 5. Conclusion

Our study proposes an approach to handle gene panel data in the context of the identification of mutations with a causal effect of the risk of phenotypes underlying a complex mode of inheritance. We visualised the approach based on data from a gene panel applied to a small cohort with BMI records available. Although the information content offered by this particular data set does not allow for accurate clinical inferences, we demonstrated the procedure of narrowing down the originally available set of variants provided by a gene panel that *per se* already consists of genes important for the phenotype under study, to SNPs that have a statistically and genomically validated potential to be causal in terms of modifying protein structure.

## Declaration of competing interest

The authors declare no competing interests.

## Ethics approval

Approval was granted by the Bioethical Committee of Poznan University of Medical Sciences (approval no. 338/1/2014).

## Consent to participate

This study was performed in accordance with the principles of the Declaration of Helsinki and consent has been obtained from each patient after full explanation of the purpose and nature of all applied procedures.

## Appendix 1

**Table A1.** Results of the SKAT test. NC represents no convergence of the estimation process. Shading highlights genes that contain significant SNPs from the mixed linear model.

| gene | chromosome | raw P values for the SKAT<br>weighted by phRED-scaled<br>SNP CADD scores | # SNPs |
| --- | --- | --- | --- |
| LEPR | 1 | 0.9169162 | 24 |
| ADIPOR1 | 1 | NC | 4 |
| AVPR1B | 1 | 1.0000000 | 7 |
| CAV3 | 3 | 0.7949824 | 17 |
| THRB | 3 | 0.2939370 | 16 |
| GHSR | 3 | 0.4777048 | 8 |
| EREG | 4 | 0.1049916 | 9 |
| NR3C2 | 4 | 0.5643788 | 9 |
| NR3C1 | 5 | 0.7322874 | 9 |
| ESR1 | 6 | 0.8102555 | 7 |
| GPER1 | 7 | 0.8590224 | 15 |
| NAMPT | 7 | NC | 5 |
| ADRB3 | 8 | NC | 3 |
| FGFR1 | 8 | 0.1870268 | 2 |
| APLNR | 11 | 0.3511696 | 10 |
| ADIPOR2 | 12 | NC | 9 |
| VDR | 12 | 0.1817712 | 18 |
| ESR2 | 14 | 0.8169512 | 5 |
| IGF1R | 15 | 0.1357984 | 33 |
| THRA | 17 | not tested | 1 |
| MC2R | 18 | 0.1112006 | 11 |
| INSR | 19 | 0.3865558 | 40 |

**Fig A1.**
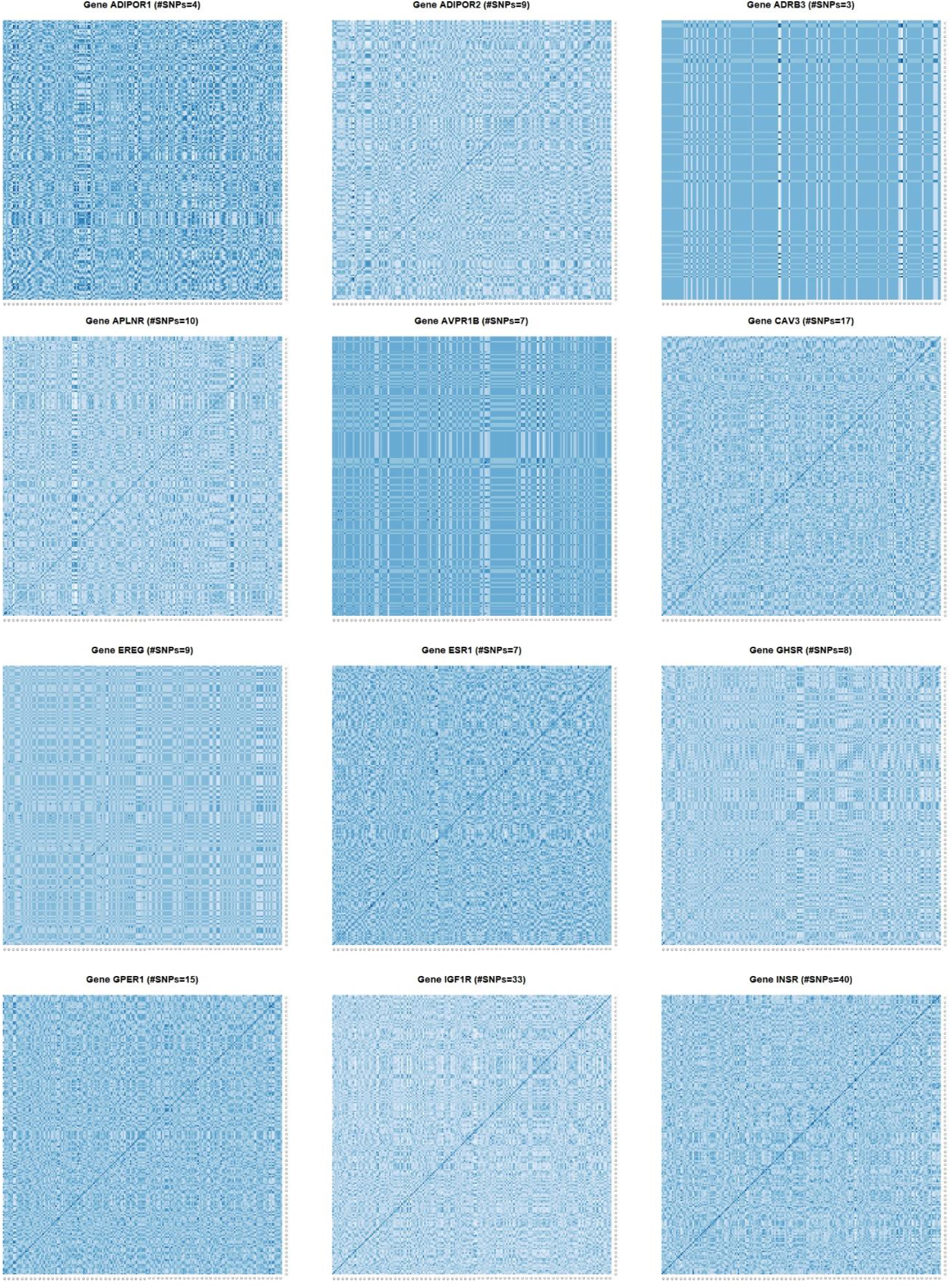
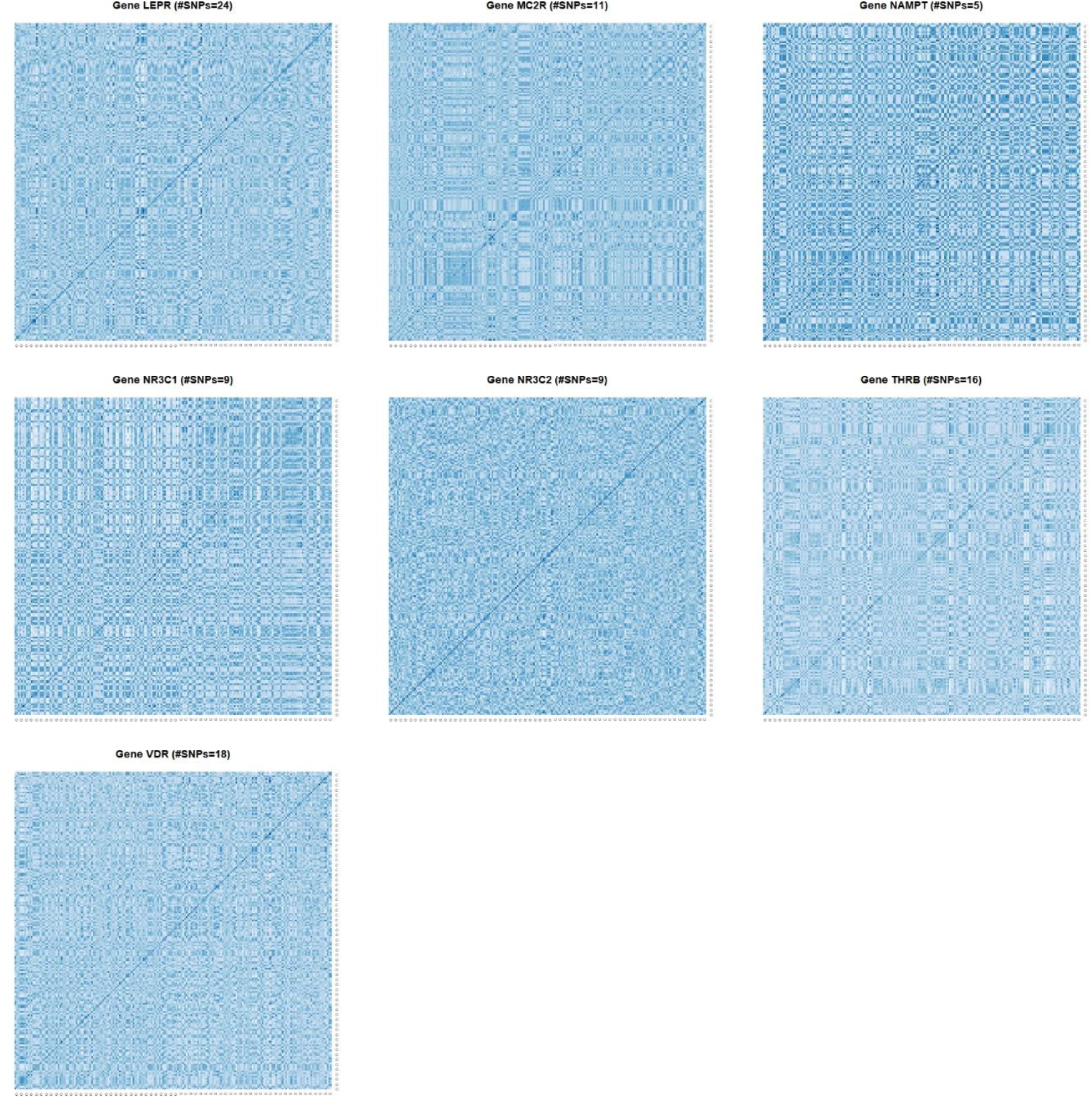
Genomic covariance matrices between individuals calculated from SNPs within each gene from the panel. Individuals were sorted according to their BMI.

## Notes

### Competing Interest Statement

The authors have declared no competing interest.

